# Seasonal phenology of Coffee Berry Borer (*Hypothenemus hampei* Ferrari) in Hawaii and the influence of weather on flight activity

**DOI:** 10.1101/2021.09.13.460180

**Authors:** Melissa A. Johnson, Nicholas C. Manoukis

**Author notes:** Corresponding author (MAJ).

## Abstract

Coffee berry borer (CBB, *Hypothenemus hampei* Ferrari) is the most serious insect pest of coffee worldwide, yet little is known about its seasonal flight behavior or the effect that weather variables have on its activity. We sampled flying female CBB adults bi-weekly over a three-year period using red funnel traps baited with an alcohol lure at 14 commercial coffee farms on Hawaii Island to characterize seasonal phenology and the influence of five weather variables on flight activity. We captured almost 5 million Scolytid beetles during the sampling period, with 81-93% of the trap catch comprised of CBB. Of the captured non-target beetles, the majority were tropical nut borer, black twig borer and a species of *Cryphalus*. Two major flight events were consistent across all three years: an initial emergence from January-April that coincided with early fruit development and a second flight during the harvest season from September-December. A linear regression showed a moderate but significant negative relationship between elevation and total trap catch. A generalized additive mixed model (GAMM) revealed that mean daily air temperature has the most significant (positive) effect on CBB flight, with most flight events occurring between 20-26 °C. Mean daily solar radiation also had a significant positive effect, while maximum daily relative humidity negatively influenced flight at values above ∼94%. Flight was positively influenced by maximum daily wind speeds up to ∼2.5 m/s and cumulative rainfall up to 100 mm, after which activity declined. Our findings provide important insight into CBB flight patterns across a highly variable landscape and will serve as a starting point for the development of flight prediction models.

## Introduction

Coffee berry borer, *Hypothenemus hampei* (Ferrari) (Coleoptera: Curculionidae) is the most damaging insect pest of coffee worldwide, causing more than $5M in annual crop losses [1]. The female coffee berry borer (CBB) initiates infestation when she bores an entrance hole into the coffee fruit (“berry”) and builds galleries for reproduction in the seed (“bean”). The offspring feed on the endosperm tissue, causing further damage to the bean and resulting in reduced quality and yields [1, 2]. Managing CBB is particularly difficult due to the cryptic nature of its life cycle which occurs almost entirely within the coffee berry. Male and female siblings mate within their natal berry, the males die, and mated females leave in search of a new berry to infest. This is the time when CBB are most vulnerable to chemical pesticides.

Strategies for managing CBB include chemical, biological, and cultural controls. Due to concerns for human and environmental health, chemical controls such as endosulfan and chlorpyrifos are being phased out or banned in many coffee-growing countries [3]. Sprays of the entomopathogenic fungus *Beauveria bassiana* are becoming more widely used as a sustainable replacement for these acutely toxic chemicals [4, 5, 6]. Biological control using parasitoids has also been implemented, particularly in Latin America, but with limited success due to the need for augmentative releases [7]. Cultural controls are typically the most cost-effective methods for managing CBB and include pruning, frequent and efficient harvesting, strip-picking all remaining berries at the end of the season, and sanitation of harvesting equipment and processing facilities [4, 5]. Integrated pest management (IPM) of CBB typically involves multiple components including sprays of *B. bassiana* early in the season, frequent and efficient harvesting, and post-harvest sanitation; these have been shown to be effective for managing CBB in Hawaii [6, 8].

A critical aspect of most successful IPM programs is monitoring of pest activity with traps and/or infestation assessments. By identifying peaks in flight activity that coincide with berry colonization, coffee growers can maximize the efficiency of *B. bassiana* applications and thereby minimize costs. Traps are used in many countries to monitor CBB activity [9, 10, 11, 12] and mass-trapping using a high density of traps in an area has even been suggested as a possible method of control [13]. In Hawaii, traps were introduced soon after the initial CBB detection in 2010 [14] as part of the IPM guidelines for managing this new invasive pest [15]. Traps were adopted by many growers in the early years of the invasion to monitor CBB activity in their fields. However, over time, fewer and fewer growers utilized traps with most transitioning to calendar sprays of *B. bassiana* or relying on casual observation of infestation in their fields to guide the timing of spray applications (A. Kawabata, pers. comm.). This approach has been shown to be inefficient: Hollingsworth et al. [6] reported that 3-5 sprays of *B. bassiana* conducted early in the season were just as effective as 8-12 calendar sprays, highlighting the importance of appropriately timing sprays for cost-effective control of CBB.

Understanding the seasonal phenology of CBB and the underlying abiotic drivers of flight are essential for predicting periods of high activity. Information on CBB population dynamics for a given coffee-growing region can be used to develop action thresholds and forecasting models for specific locations. This is especially important for Hawaii where the coffee-growing landscape is heterogeneous, and a single set of management recommendations is difficult to apply across the entire region. Coffee is grown commercially on six of the main Hawaiian Islands on volcanic soils that vary in age and nutrient composition and experience a broad range of microclimate conditions from sea level to over 800 m in elevation [16]. On Hawaii Island, more than 800 small farms produce coffee, most of which are family-run operations that rely on manual labor to harvest the coffee and implement management practices. Optimizing pest management strategies while minimizing costs are critical to the longevity of these farms as Hawaii has some of the highest labor and production costs of any coffee-growing region.

In the present study we monitored the flight activity of CBB using funnel traps at 14 commercial coffee farms in the two main coffee-growing regions of Hawaii Island, Kona and Kaʻu, over a three-year period. Five weather variables thought to be important for insect flight (temperature, relative humidity, solar radiation, wind speed and rainfall) were tracked at each site. Our objective was to provide insights into the seasonal phenology of CBB across a highly variable landscape, as well as to characterize the abiotic factors that trigger CBB flight, which will aid in the development of models for predicting future flight events. These models can serve as decision support tools to guide the timing of pesticide applications. While this study is focused on Hawaii, we expect that the information gained on CBB flight and associated weather variables will be useful for developing IPM strategies in other coffee-growing regions.

## Materials and Methods

### Study Sites

Fourteen commercial coffee farms were selected for the study on Hawaii Island. Eight farms were located on the West side of the island in the Kona district and six farms were on the Southeast side of the island in the Kaʻu district (Fig. 1). While coffee is grown throughout the Hawaiian Islands, Kona and Kaʻu are the two primary coffee-growing regions, both of which are world-renowned for the high quality of their coffee. Farms were selected to encompass the broad range of elevations and climatic conditions under which coffee is grown on Hawaii Island. In Kona, farms ranged in elevation from 204 m – 607 m and in Kaʻu farms ranged from 279 m – 778 m in elevation. All farms were actively managed for coffee production throughout the study, although the timing and number of interventions varied. Management practices included regular pruning, weed management, fertilizer, pesticide application, cherry harvesting and end of season strip-picking. Farms were largely characterized as sun-grown although some farms had scattered fruit, nut or ornametal shade trees planted as well. All farms had *Coffea arabica* var. *typica* planted with the exception of one farm in Kaʻu which had primarily var. *catuai* planted.

**Figure 1.**
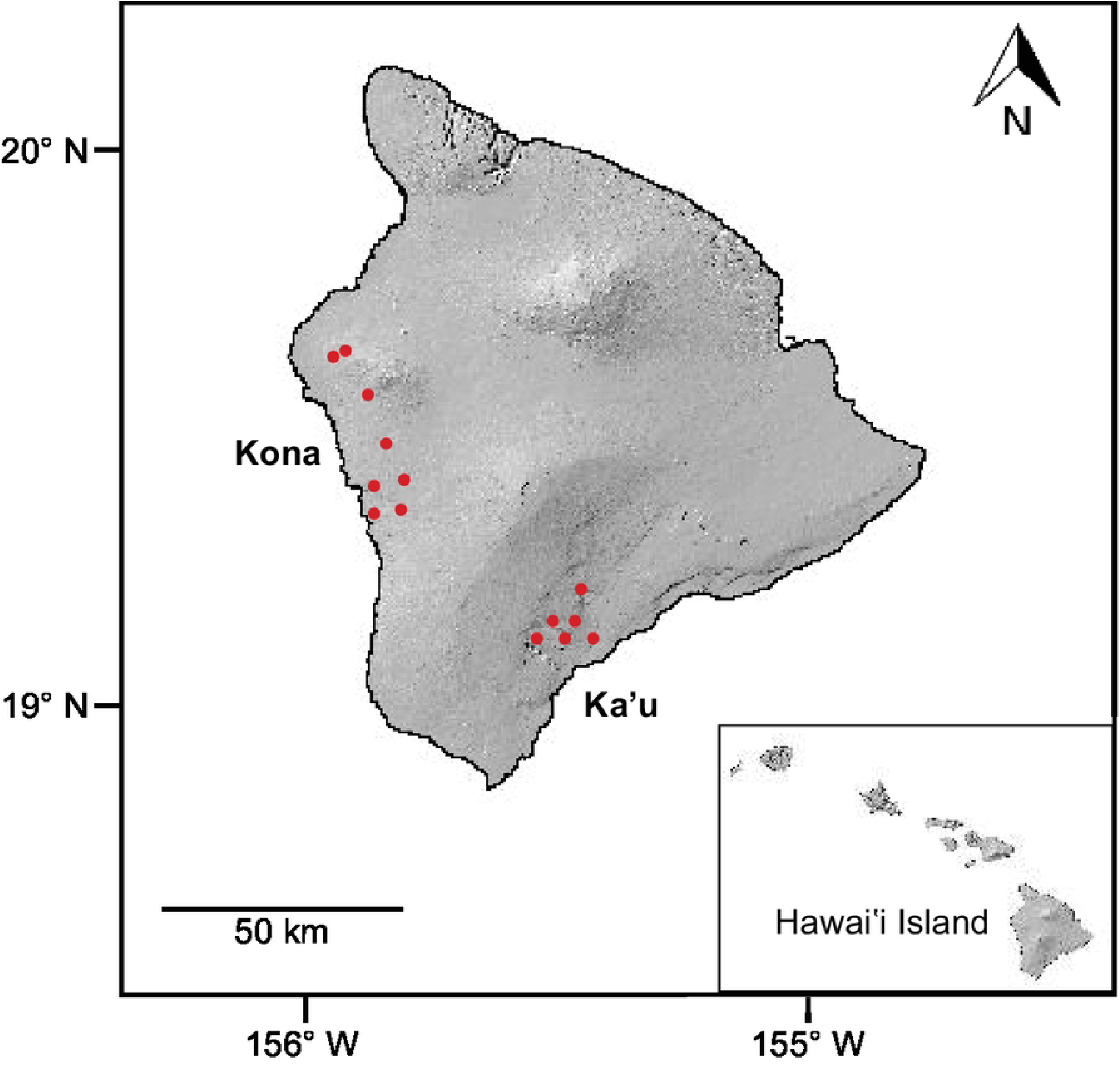
Map of Hawaii Island showing 14 study sites (eight in the Kona district and 6 in the Kaʻu district). Inset map shows the main Hawaiian Islands and the location of Hawaii Island within the archipelago.

### Flight Activity

Red funnel traps (CIRAD, Montpellier, France) baited with an alcohol lure (3:1 methanol:ethanol) were randomly distributed throughout each farm. Trap density was based on farm size, with 3-5 traps used for small farms (1-1.4 ha) and 6-9 traps used for large farms (1.5-2 ha). Traps were hung on stakes at ∼1 m in height and were equipped with a collection cup containing propylene glycol. Trap contents were collected in 70% ethanol on a bi-weekly schedule from 2016-2018. Lures were refilled as needed and propylene glycol was replaced bi-weekly. In the laboratory, trap contents were passed through a sieve (1.5 mm mesh size) to seperate out all large insects, which were discarded (see [17] for additional details on trap setup, collection and processing). The remaining insects from each trap collection were placed under a stereomicroscope (Leica microsystems GmbH, Wetzlar, Germany). All Scolytine beetles were counted and CBB were seperated from these other beetles to estimate trap specificity. If >500 beetles were caught in a single trap, we used a volumetric method to estimate count (see [17] for details). For each site, the number of CBB per trap per day (CBB/trap/day) was estimated by dividing the total number of CBB caught by the number of traps in the farm and then dividing this by the number of days in the sampling period (14 days on average, although this varied occasionally). This calculation was also done to estimate the number of other Scolytid beetles caught per trap per day.

### Weather Variables

Manual or cell-service weather stations were set up at each farm to measure the following variables: air temperature, relative humidity (RH), rainfall, wind speed and solar radiation. Manual stations consisted of a Hobo Pro v2 temperature/RH data logger (U23-002, Onset Computer Corporation, Bourne MA) housed in a solar shield (RS3, Onset Computer Corporation, Bourne MA), a solar pendant (UA-002-64, Onset Computer Corporation, Bourne MA) and a rain gauge equipped with a manual data logger (RainLog 2.0, RainWise Inc.). Cell-service weather stations were comprised of a 4G remote monitoring station (RX3004-00-01, Onset Computer Corporation, Bourne MA) equipped with a temperature/RH sensor (S-THB-M002), solar panel (SOLAR-5W), solar radiation sensor (S-LIB-M003) placed within a solar shield (RS3-B) and rain gauge (S-RGB-M002). Wind speed sensors (S-WSET-B) were added to each cell-service station in 2018. For each site, the daily maximum, mean, and minimum were estimated for air temperature and RH in R v. 3.5.0 using the ‘aggregate’ function in the *stats* package [18]. We used the same method to estimate the daily mean and maximum wind speed and solar radiation, as well as daily cumulative rainfall. We then used these daily values to calculate the average air temperature (°C), RH (%), wind speed (m/s) and solar radiation (W m^-2^), as well as the cumulative rainfall (mm) for each ∼bi-weekly sampling period.

### Data Analysis

All statistical analyses were conducted in R v.3.5.0 [18]. The assumption of normality for each variable was validated using quantile-quantile plots and a Shapiro-Wilks test; an *F* test was conducted to assess for equal variances using the *stats* package. A Pearson correlation test was conducted using the *stats* package to examine the relationship between total CBB capture for each year/site and elevation. The mean number of CBB caught per trap per day was log-transformed (log + 1) prior to analysis. Linearity of the relationship between CBB/trap/day and each weather variable was also checked prior to analysis using two-dimensional scatterplots. Given the non-linear nature of the relationship between weather variables and CBB flight, the influence of weather on CBB flight activity was evaluated with a generalized additive mixed model (GAMM) in the *mgcv* package v. 1.8-23 [19].

The response variable for the model was the log-transformed mean number of CBB/trap/day, with year and site included as random effects and the weather variables included as fixed effects with cubic regression splines. We assumed a gaussian error distribution and an identity link function for the model. We used the generalized cross-validation (GCV) score to measure model smoothness with respect to the smoothing parameters as well as the estimate prediction error. A lower GCV score indicates a smoother model and is somewhat comparable to an Akaike Information Criterion (AIC) value, in that a lower score equates to a better fitting model. Pairwise Pearson correlation tests of continuous explanatory variables were conducted to assess multicollinearity at a correlation coefficient threshold of 0.7; maximum temperature, mean RH and maximum solar radiation were subsequently dropped from the model due strong correlations with mean temperature, maximum RH and mean solar radiation, respectively.

## Results

In total, just under 5 million Scolytid beetles were captured over a period of 143 weeks. Across all 14 sites, CBB made up an average of 81% of the total trap catch in 2016, 91% in 2017, and 93% in 2018 (Fig. 2). The most trapped non-CBB beetles were *Hypothenemus obscurus* F. (tropical nut borer), *Xylosandrus compactus* Eich. (black twig borer) and a species of bark beetle tentatively identified to the genus *Cryphalus* Erichson. Peak activity for non-CBB beetles was in the summer months of June and July. Farms observed to have higher percentages of non-CBB beetles in traps were located next to macadamia nut orchards or forests or had other fruit and nut trees interplanted with the coffee. The combined CBB catch across all sites was similar for 2017 (∼1.98M CBB) and 2018 (∼2.05M CBB), but considerably lower for 2016 (∼600K CBB) given that data collection did not start until March (Kona) and May (Kaʻu), thereby missing the initial emergence for that year.

**Figure 2.**
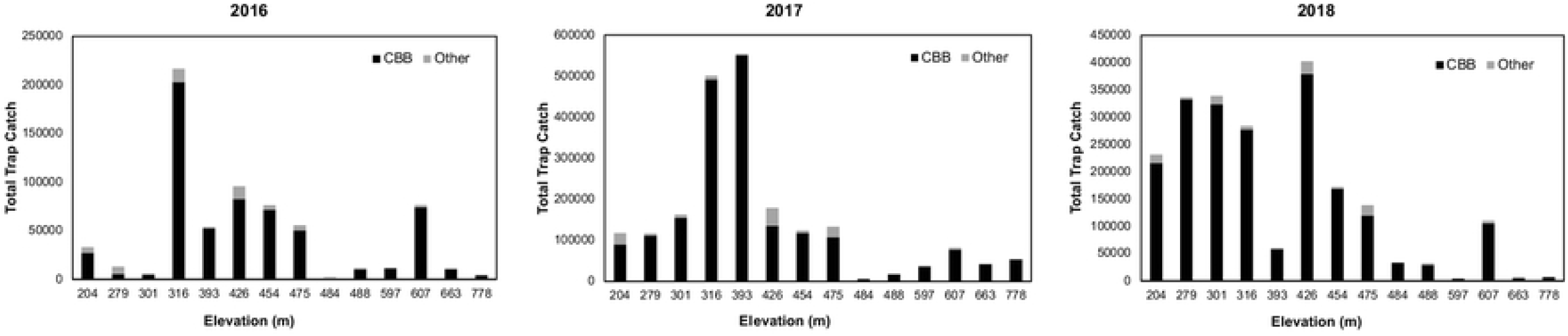
Total trap catch across three years on Hawaii Island. Study sites are in order of increasing elevation from left to right. Black bars represent coffee berry borer (CBB) while gray bars represent other Scolytid beetles.

Although variation was observed among years and farms in terms of the average number of CBB/trap/day, the general pattern of flight was consistent (Fig. 3). Seasonal phenology was observed in two stages: an initial emergence from January-April which coincides with early fruit development, and a secondary flight which occurs from September-December and coincides with the harvest season (Fig. 3). This secondary flight corresponds to the emergence of new generations of CBB that were the offspring of the initial colonizing females. Although we did not observe any consistent differences in trap capture patterns among farms in the Kona vs. Kaʻu regions, a linear regression revealed a moderate but significant negative correlation between elevation and total trap catch (*R* = -0.48, t = -3.39, p = 0.002) (Fig. 4).

**Figure 3.**
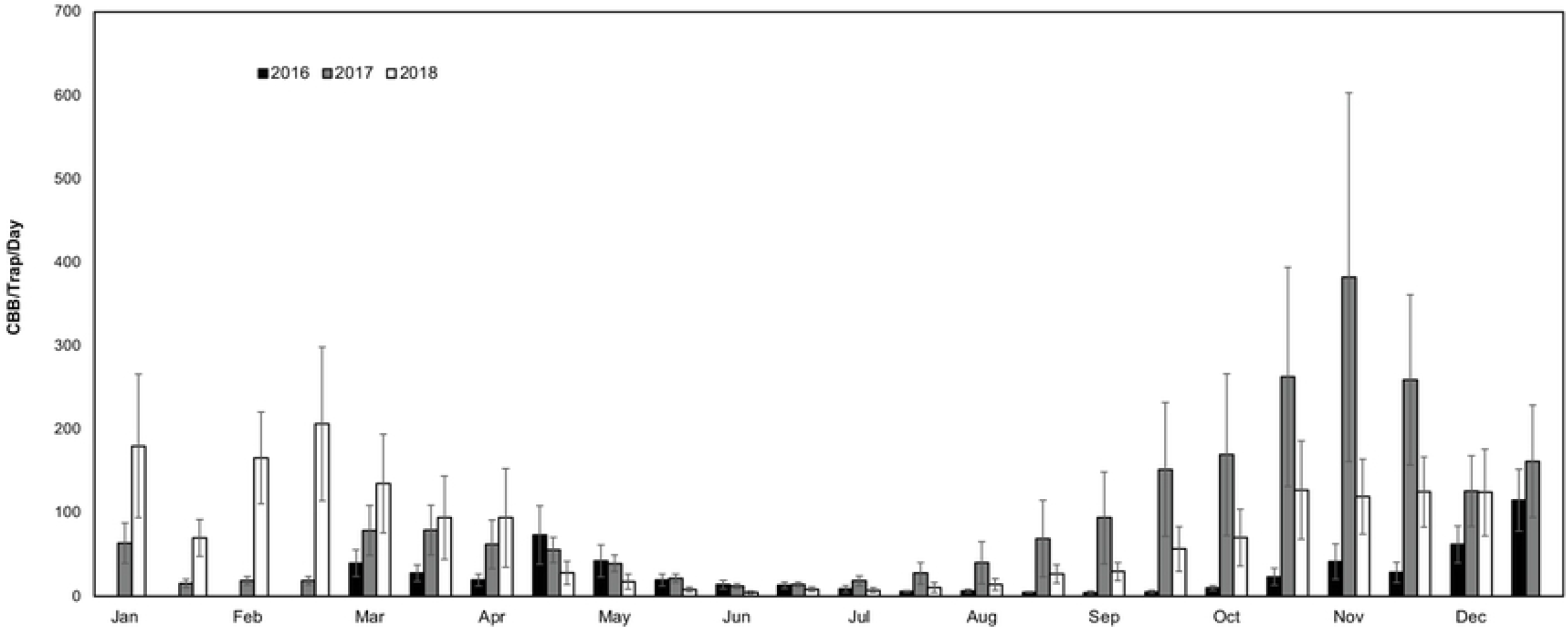
Seasonal flight phenology of coffee berry borer (CBB) on Hawaii Island over a three-year period. Sampling began in March 2016 and ended in December 2018. Error bars show the variation across 14 study sites.

**Figure 4.**
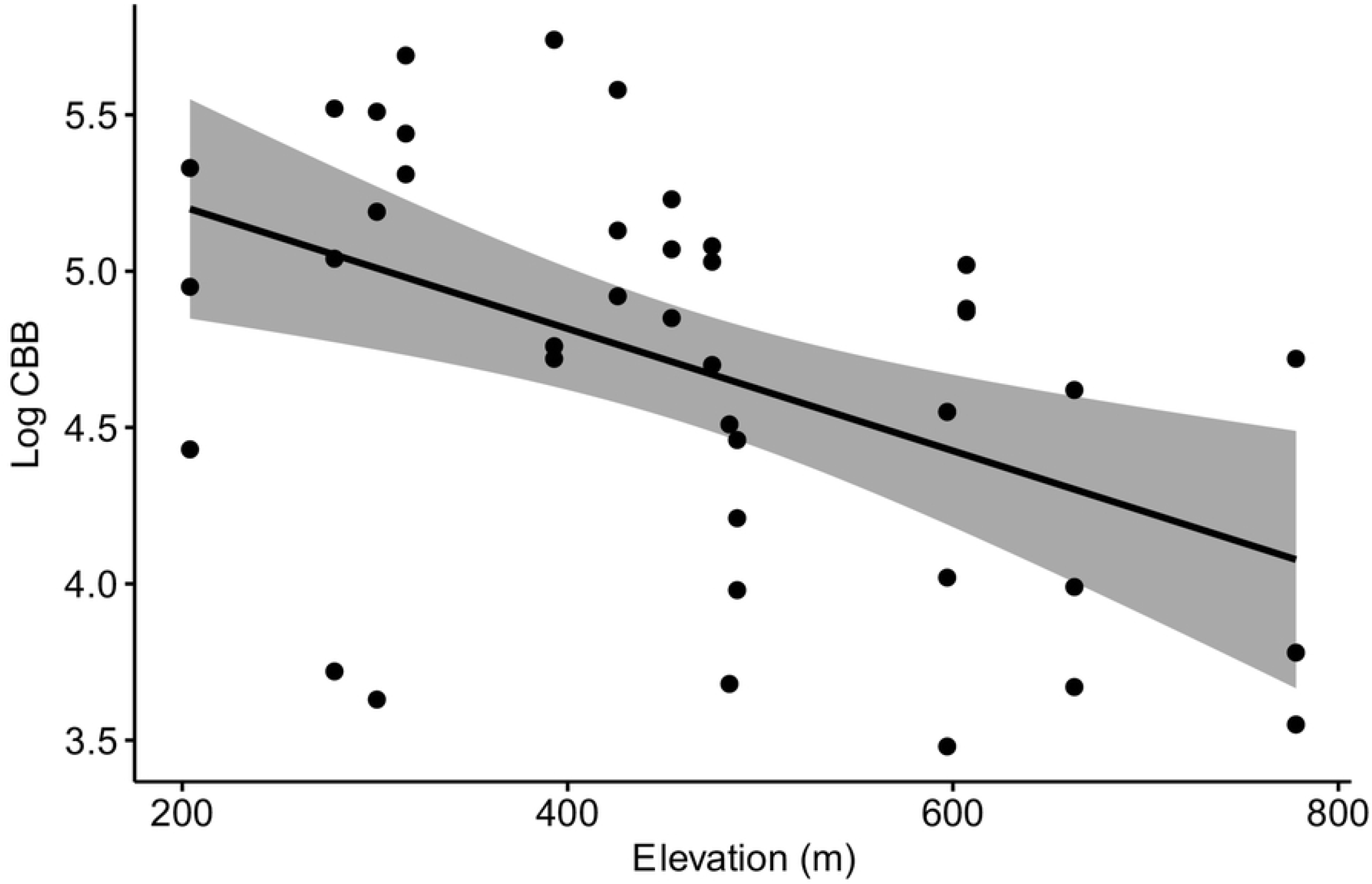
Linear regression showing a moderate but significant negative relationship between the log-transformed total CBB capture for each site/year and elevation (R = -0.48, p = 0.002).

The final GAMM explained 75.7% of the deviance (adjusted R^2^ = 0.66, GCV = 0.29), with the following weather variables having a significant effect on CBB flight at an alpha < 0.05: mean temperature, maximum RH, cumulative rainfall, maximum wind, and mean solar radiation (Table 1). Mean daily temperature had the greatest positive effect on CBB flight (Table 1) with three peaks observed between 20-26 °C (Fig. 5A). Mean daily solar radiation also had a positive effect on CBB flight, with peaks observed at 200-300 W/m^-2^ and 400-500 W/m^-2^ (Fig. 5A). Flight levels were generally high at maximum daily RH values between 80-94%, after which they fell sharply (Fig. 5B). Cumulative rainfall had a positive effect on CBB flight up to 100 mm, after which flight decreased (Fig. 5B). Flight increased up to maximum daily wind speeds of ∼2.5 m/s and subsequently dropped off (Fig. 5C). Both the random effects of site (p < 0.001) and year (p = 0.003) also had significant effects on CBB flight (Table 1).

**Table 1.**
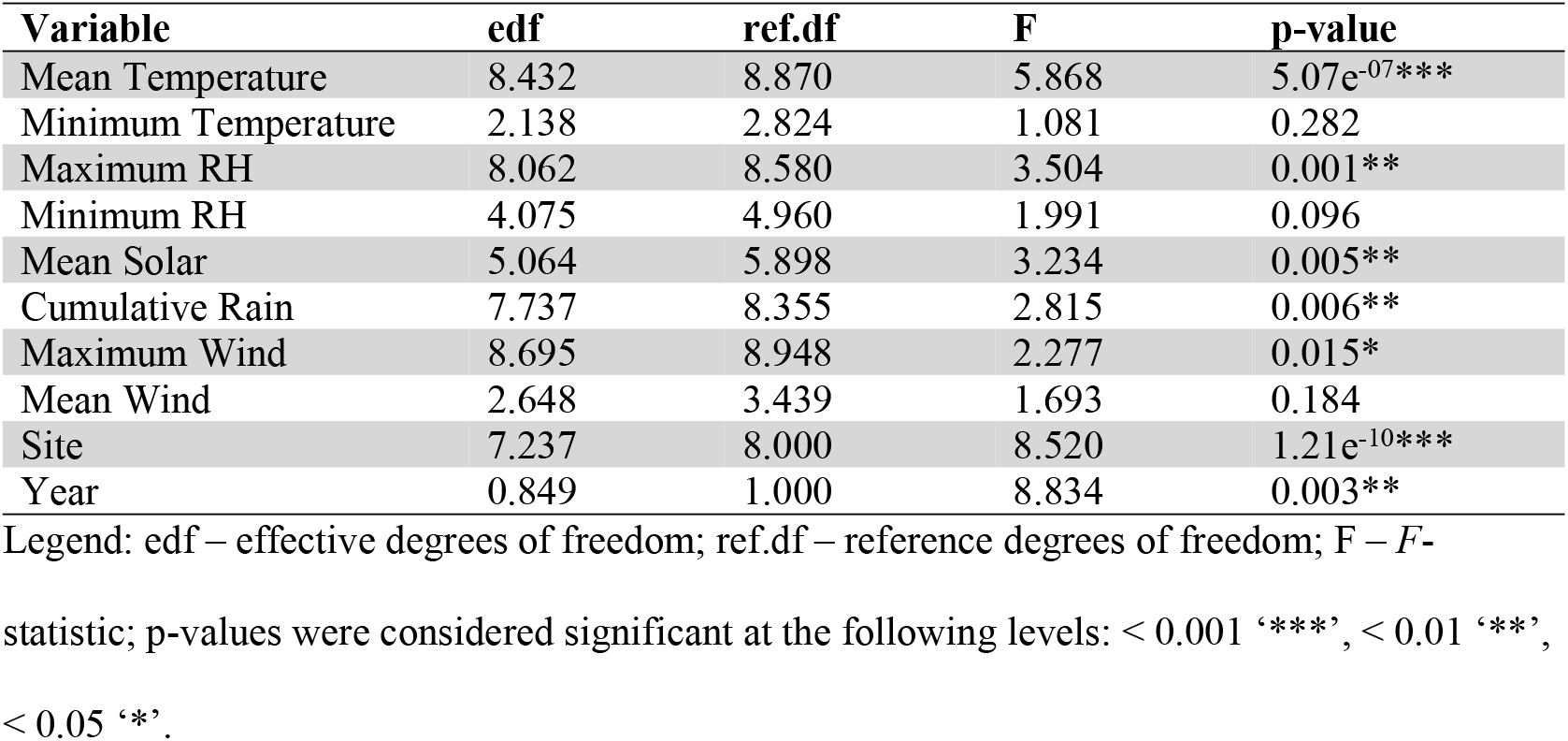
Results from the generalized additive mixed model (GAMM) analyzing the influence of five independent weather variables (fixed effects) and site and year (random effects) on CBB flight.

| Variable | edf | ref.df | F | p-value |
| --- | --- | --- | --- | --- |
| Mean Temperature | 8.432 | 8.870 | 5.868 | 5.07e<sup>-07***</sup> |
| Minimum Temperature | 2.138 | 2.824 | 1.081 | 0.282 |
| Maximum RH | 8.062 | 8.580 | 3.504 | 0.001** |
| Minimum RH | 4.075 | 4.960 | 1.991 | 0.096 |
| Mean Solar | 5.064 | 5.898 | 3.234 | 0.005** |
| Cumulative Rain | 7.737 | 8.355 | 2.815 | 0.006** |
| Maximum Wind | 8.695 | 8.948 | 2.277 | 0.015* |
| Mean Wind | 2.648 | 3.439 | 1.693 | 0.184 |
| Site | 7.237 | 8.000 | 8.520 | 1.21e<sup>-10***</sup> |
| Year | 0.849 | 1.000 | 8.834 | 0.003** |

Legend: edf – effective degrees of freedom; ref.df – reference degrees of freedom; F – *F*-statistic; p-values were considered significant at the following levels: < 0.001 ‘\*\*\*’, < 0.01 ‘\*\*’, < 0.05 ‘\*’.

**Figure 5.**
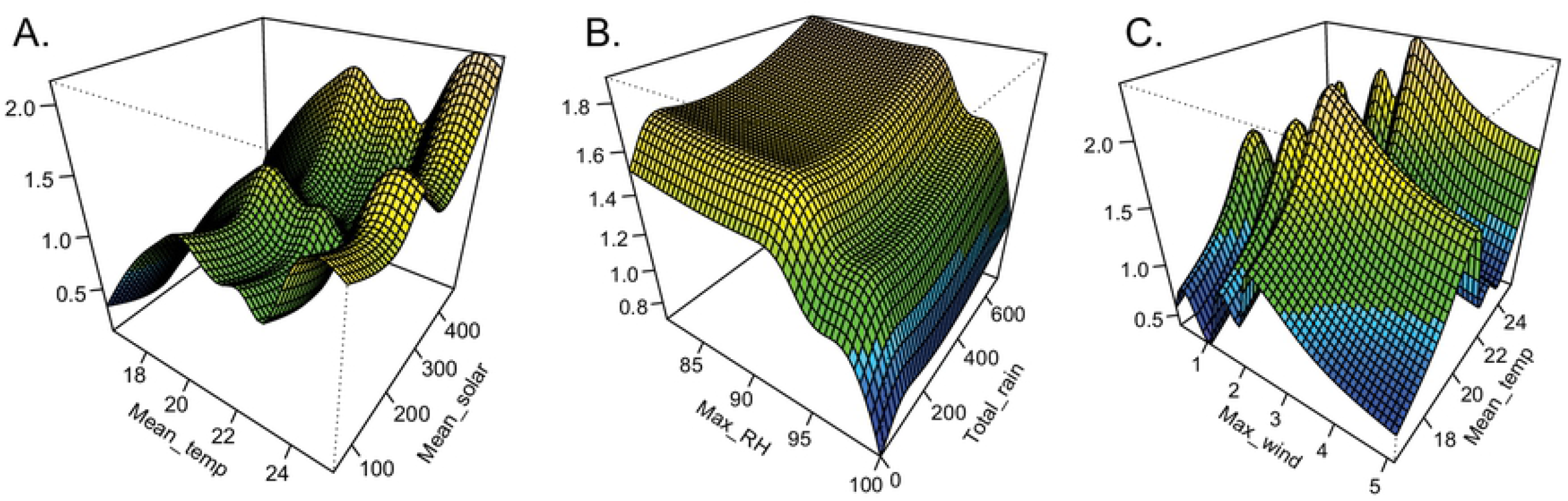
Results from a generalized additive mixed model (GAMM) exploring the effects of mean daily temperature (A, C), mean solar radiation (A), maximum relative humidity (B), cumulative rainfall (B) and maximum wind speed (C) on coffee berry borer flight activity. Three-dimensional contour plots show peaks in CBB flight activity in yellow and decreased activity in cooler colors (green and blue).

## Discussion

Determining the seasonal phenology and abiotic factors involved in flight activity is a critical step in developing an integrated pest management plan for invasive insects. In the present study we examined 14 commercial coffee farms in two coffee-growing districts on Hawaii Island to elucidate seasonal flight patterns and the influence of five weather variables on CBB flight. Our findings suggest that although there are differences from farm to farm, general patterns of CBB flight activity can be described across this highly variable landscape. We observed two major flight events that were consistent across all three years: an initial emergence from January-April that coincides with early fruit development and a secondary flight that occurs during the harvest season from September-December. We also found that despite not having a species-specific lure, trap specificity was generally high with CBB making up 81-93% of the total trap catch across all farms.

These results correspond to findings from two earlier studies that examined CBB flight patterns on Hawaii Island. Messing [20] reported CBB flight from November-March at two farms in Kona. That study also found that 1:1 and 3:1 ratios of methanol:ethanol captured similar numbers of beetles, and that non-target beetles made up an insignificant proportion of trap catch relative to CBB (3-7 non-target beetles/trap/day compared to 100-400 CBB/trap/day). Our findings are also in line with those of Aristizábal et al. [21], who examined flight activity at 15 farms in Kona and Kaʻu using 3:1 methanol:ethanol baited funnel traps. That study reported a small peak in flight from May-July and a larger peak from December-February. The authors suggested that peaks appeared to coincide with increased levels of rainfall following a dry period, although weather data was not available for each farm, excluding statistical analyses. Aristizábal et al. [21] also estimated that non-target insects made up < 5% of all trap catch based on subsampling from a few farms. While these earlier studies provided initial insights into seasonal flight trends in Hawaii, sampling was limited to a one-year period and did not include site-specific data on weather. In the present study we were able to expand on these initial findings by collecting data over multiple years and correlating flight activity to five individual weather variables at each farm. Below we summarize our main findings with respect to the influence of weather variables on trap catch, our proxy for flight activity.

Mean daily air temperature was observed to be the single weather variable with the strongest (positive) relationship to CBB flight activity across all sites. This is relatively unsurprising since insects are poikilotherms, meaning their body temperature depends on ambient environmental temperature. Many aspects of insect biology are driven by temperature including generation length, rate of development, mating activity and dispersal [22]. It is widely reported that insects actively regulate body temperature before and during flight by behavioral or physiological means [23; 24; 25; 26; 27; 28; 29], and that there is a lower and upper temperature threshold for flight [30]. Insects need warm temperatures to initiate and maintain flight as it becomes difficult at lower temperatures to generate heat [27, 31, 32, 33]. As air temperatures increase, the aerodynamic force and mechanical power output of insects is also increased along with wing beat frequency [34, 35].

Chen and Seybold [36] reported a lower threshold of 11 °C, an optimum of 27 °C and an upper threshold of 39 °C for the walnut twig beetle (*Pityophthorus juglandis* Blackman). In a controlled laboratory setting with RH held at 90% and 100%, Baker et al. [37] reported low CBB emergence from dried berries at temperatures below 20 °C, a marked increase in emergence from 20-25 °C, and no significant increase above 25 °C. In the present study under highly variable field conditions, we observed that most flight events took place when mean daily temperatures were between 20-26 °C, with very few events below 16 °C or above 32 °C.

Along with a positive effect of increasing temperature, we found a positive significant relationship between CBB flight and mean daily solar radiation. This is line with the findings of several studies that reported temperature and solar radiation as the main abiotic factors positively influencing beetle flight [36, 38, 39, 40]. Related to this, we also observed a significant negative correlation between elevation and total CBB capture. Hamilton et al. [41] showed that CBB development on Hawaii Island is faster at low elevations primarily due to higher temperatures at these locations. The authors estimated 4-5 generations per season at low elevations (200-300 m), compared to 2-3 generations per season at high elevations (600-800 m). Thus, the higher abundance of CBB caught at lower elevations is directly related to the shorter development times at these sites.

In accordance with numerous studies on insect flight, we observed a significant negative effect of maximum daily relative humidity above ∼94%. CBB may be reluctant to leave the berries during periods of very high RH as this may indicate rain (along with an associated drop in barometric pressure, not measured in the current study), causing them to shelter. In addition, greater CBB mortality can occur during periods of high RH due to proliferation of *B. bassiana* under moist, humid conditions. Lastly, at very high RH there is a higher requirement of wing-beat frequency, which is metabolically costly [23, 29]. Farnworth [42] showed that *Periplaneta americana* Linnaeus had a higher wing beat frequency at 95% RH compared to 50% RH at temperatures between 27-35 °C, which could reflect greater effort to dissipate heat at high vs. low humidity. In contrast to our findings, under laboratory conditions with temperature held at 25 °C, Baker et al. [37] described high CBB emergence from infested two-month-old berries at RH values of 20% and 55%, minimum emergence at 78% and 90%, and a steady increase in emergence from 94-100%. It is likely that differences between this study and our findings are related to the setting under which emergence was estimated. We did not observe a minimum RH lower than 52%, and mean RH values ranged from 80-90% for most sites/years. Maximum RH values were typically >94% through spring and summer and dropped during the winter months (November-March), which coincided with peak CBB flight.

We observed a similar trend for maximum wind speed, with flight increasing at wind speeds up to ∼2.5 m/s and then decreasing at speeds above that value. Chen and Seybold [36] reported that *P. juglandis* flight was limited at very low wind speeds and peaked when temperature was ∼30 °C and wind speed was 2 km/h. Pawson et al. [43] reported low flight activity of the bark beetle *Hylurgus ligniperda* Fabricius at very low wind speeds, an increase with rising wind speeds, and a peak at 2 m/s. Thus, some wind may help to initiate flight as well as provide olfactory cues to allow detection of resources, but at very high wind speeds flight appears to be inhibited. The long-distance dispersal of many weak-flying smaller insects is dictated mostly by wind [44, 45, 46], which may explain the tendency of CBB to fly near the ground (M. Johnson, unpub. data) where ambient wind speeds are generally low [47]. Lastly, cumulative rainfall had a positive effect on CBB flight up to a point; flight appeared to be inhibited during periods of heavy rainfall (>100 mm). Other studies have reported negative effects of heavy rainfall on bark beetle flight [48, 49, 50]; the very small size of CBB likely precludes its movement during periods of inundation.

By considering the phenology of the coffee crop and the CBB together with the trap catches reported here, the pattern of CBB movement in Hawaii coffee plantations comes into focus. At the very beginning of the year the new coffee crop must become mature enough (>20% dry matter content [51, 52, 53]) to be infested by the previous season’s beetles. These CBB are mostly in raisins (dried berries) on the ground under the coffee trees or on the raisins remaining on the branches-the latter being the most heavily infested repositories on a per-bean basis [54]. The first flight suggests movement from these refugia into developing green berries in the first quarter of the calendar year. Given that few berries may be available early in the season, CBB will have to travel more extensively to find suitable hosts. Once a suitable berry is located, the female CBB will penetrate the exocarp and wait to complete entry into the coffee bean until conditions are suitable, or may begin boring immediately into the seed depending on fruit stage [1].

After the first flight, CBB begin reproducing within the beans and berry development continues. During this middle part of the year, there is very little baseline movement of CBB and trap catches are low since the host berries are abundant. The second major flight, observed here in the last quarter of the calendar year, is likely driven by waning food supplies in the original berries (as the numbers of CBB increase in the individual beans, food is reduced and movement again becomes necessary) in combination with physical disturbance during harvesting, strip-picking and tree pruning. Based on observations of lower flight activity in this second part of the year in feral and unmanaged sites vs. well-managed sites on Hawaii Island [55], physical disturbance via management practices are a larger stimulus for this second flight relative to the need to locate food, which is the primary stimulus for the first flight of the season.

## Conclusions

Our investigation into CBB seasonal phenology and the influence of weather on CBB flight activity revealed important insights that will be useful for the development of future flight prediction models. Seven of the 10 variables included in the GAMM were found to have a significant influence on CBB flight activity. Of the eight fixed effects, mean daily temperature, mean solar radiation, maximum RH, maximum wind speed and cumulative rainfall were significant, while both the random effects of site and year also had a significant effect on flight. Variation among years was likely due to a combination of weather events and the timing of study initiation, while variation among farms likely involved differences in management style, timing, and the number of interventions, as well as differences in CBB development times across elevations. That the best model explained only 75.7% of the observed variance likely reflects the absence of additional weather variables that were not measured such as barometric pressure, as well as variation that was missed due to sampling frequency. Given the large travel distance between sites, all traps were sampled on a bi-weekly schedule, such that correlating weather events to trap catch at finer scales was not possible. The timing of daily CBB flight and the weather events correlated with flight activity on an hourly basis are currently being examined at several sites on Hawaii Island to further elucidate flight patterns in this global pest of coffee.

## Acknowledgements

Robbie Hollingsworth, Ray Carruthers and Luis Aristizábal made helpful suggestions on study design and methods. Many technical staff steadfastly collected the data required for this study, regularly under difficult conditions. These included Matthew Mueller, Thomas Mangine, Samuel Fortna, Austin Bloch, Jaqueline Pitts, John Ross, Lori Carvalho, Shannon Wilson, Colby Maeda, Connor Rhyno and others. Special thanks to Forest Bremer for developing the electronic data collection system for our monitoring program. We are also grateful to the Kona and Kaʻu coffee growers that allowed us to conduct this study on their farms. Opinions, findings, conclusions, or recommendations expressed in this publication are those of the authors and do not necessarily reflect the views of the USDA. The USDA is an equal opportunity provider and employer.

